# Use of the Pacific oyster (*Magallana gigas*) as a natural sampler for the detection of *Neoparamoeba perurans*

**DOI:** 10.1101/2025.08.06.669027

**Authors:** Brett Bolte, Andrew Bissett, Pascal Craw, Carmel McDougall, James W. Wynne

**Affiliations:** CSIRO Environment, Hobart, TAS, Australia; Blue Economy Cooperative Research Centre, PO Box 897, Launceston, TAS, Australia; Griffith University School of Environment and Science, Nathan, QLD, Australia; Australian Rivers Institute, Nathan, Australia; CSIRO Agriculture and Food, Hobart, Australia; Scottish Oceans Institute, University of St. Andrews, St. Andrews, Scotland

**Keywords:** AGD, natural samplers, eDNA

## Abstract

Amoebic gill disease (AGD), caused by *Neoparamoeba perurans,* is a challenge for Atlantic salmon aquaculture. Research has, therefore, focused on detecting/monitoring *N. perurans* loads within/around fish pens. Recently, molecular methods to detect *N. perurans* have been used to reduce labour-intensive sampling, inconsistency, and stress on stock, while complimenting gill scoring and histology methods for AGD assessment. Molecular detection depends on reliable and simple sample collection, opening the possibility of using natural samplers (organisms that collect/accumulate DNA through feeding/foraging) for aquatic eDNA collection. We evaluated, through aquarium-based experiments, the utility of the Pacific oyster (*Magallana gigas*) to collect *N. perurans* DNA from the water column. In aquaria inoculated with *N. perurans*, total water column amoeba load decreased significantly over time in the presence of oysters. Despite this decrease no correlation between the level of amoeba accumulation within oysters and the decrease in the water column was observed. *N. perurans* was detected in every oyster organ type tested (mantle, gill, palp and digestive gland), though with high variation. The detection of *N. perurans* DNA within the digestive gland indicates that oysters ingested amoeba. Oysters were found to be viable eDNA samplers for *N. perurans*, providing a useful, simplified collection method.

## 1. Introduction

*Neoparamoeba perurans* is an economically important protozoan parasite and is the causative agent of amoebic gill disease (AGD). The disease has been reported in 15 finfish species (Han, 2019) and is of great concern in finfish aquaculture, especially within Atlantic salmon (*Salmo salar*) farming. *N*. *perurans* infection causes gill lesions (Bridle et al., 2010, Han, 2019), compromising the gills through the decline in the functional surface area available for gas and ion exchange (Hvas et al., 2017). If left untreated it can cause mortality rates of 50 – 80 % (Adams and Nowak, 2003, Oldham et al., 2016, Talbot et al., 2022). Initially discovered to be affecting Tasmanian pen-farmed Atlantic salmon in the 1980s (Munday, 1986), AGD prevalence in aquaculture has increased globally (Steinum et al., 2008, Bustos et al., 2011). On Atlantic salmon farms, routine monitoring of AGD is typically performed by gross gill scoring of the severity of presumptive AGD gill pathology (Taylor et al., 2009), which is a low cost, but labour intensive, real-time method. Histopathology is considered the reference standard AGD diagnostic, as it confirms the presence of amoebae associated with gill lesions (Adams et al., 2004). Finally, gill swab qPCR is used to confirm that the observed amoeba is *N. perurans* (Downes et al., 2017). Both histology and qPCR require the same labour-intensive sampling as gill scoring but incur additional costs and may be slow to support husbandry decisions. While husbandry decisions may be delayed, qPCR enables quantitative assessments of amoeba load within the sample. Understanding the relationship between amoeba loads within the water and disease onset/progression is of interest due to its potential to provide rapid, non-invasive detection, warning of impending disease outbreaks (Bridle et al., 2021). To determine water column amoeba loads, amoeba (or its DNA) must be successfully collected from the environment.

Environmental DNA (eDNA) methodologies provide sensitive detection of a target(s) via amplification of (trace) amounts of DNA left behind in the environment (Turner et al., 2015, Cavallo et al., 2018, Walsh et al., 2018). For this study, we use the definition of eDNA that includes both trace DNA and DNA contained within whole microbes (Lugg et al., 2018). A successful eDNA campaign aims to collect sufficient DNA for downstream analyses, collect DNA in an efficient and/or cost-effective manner, and protect/preserve DNA. The most commonly used methods in aquatic environments typically either utilise pump-based filtration of water through a membrane to collect DNA and particulates, or precipitation/centrifugation of DNA directly from water samples (Hinlo et al., 2017).

Filtration through a variety of filter sizes (0.22 µm, 0.45 µm) and centrifugation have successfully been used for the collection of *N. perurans* DNA from the water column (Bridle et al., 2010, Wright et al., 2015, Wright et al., 2017, Taylor et al., 2021, Wynne et al., 2024) providing effective methods to collect *N. perurans* DNA from the water column while minimising stock stress.

Recent studies have demonstrated that many organisms naturally collect DNA from the environment through feeding/foraging, and that these organisms can be used as ‘natural samplers’ for eDNA collection (Schnell et al., 2015, Siegenthaler et al., 2019, Drinkwater et al., 2019, Mariani et al., 2019, Turon et al., 2020, Cai et al., 2022, Weber et al., 2021, Jeunen et al., 2023). Filter feeding organisms are often co-located with Atlantic salmon farms, where salmon farm effluent has a positive impact on their growth (Byrne et al., 2018). Utilising these organisms as natural samplers would negate the need for pump-based water filtration, potentially enabling a simplified eDNA collection method.

Oysters are common in marine environments, including around fish farms, and have been shown to consume/ingest a wide range of particles from 4-72 µm in size (Dupuy et al., 1999). *Neoparamoebae* fall within this size range (10-20 µm; (MacPhail et al., 2021) and thus may also be successfully filtered from the water column. To date, two studies have looked at the ability of bivalves to accumulate *N. perurans* (Hellebø et al., 2017, Rolin et al., 2016). Hellebø and colleagues (2017) detected *N. perurans* within a small percentage (5/49 replicates) of blue mussel (*Mytilus edulis*) gill or digestive gland samples. These samples were taken from a single fish pen where amoebae were detected in the water column (determined by qPCR ct values), and in which farmed fish had severe AGD affected gills. In a lab-based study, *N. perurans* inoculated in high concentrations (3000 cells per L^-1^) was not detected in blue mussel organs (pooled gill and mantle), but was detected from shell swabs (Rolin et al., 2016). Results from both studies suggest that blue mussels are poor natural samplers for *N. perurans,* highlighting the need to explore other bivalves that may be better suited.

Different filter feeding species, even within the ostreid family, have great variability in their particle retention (Nielsen et al., 2016). Therefore, it is possible that Pacific oysters (*Magallana gigas*) may have better capacity than blue mussels for accumulation of amoeba DNA. This study investigated the ability of *M. gigas* to accumulate *N*. *perurans* (as detected via a DNA based assay) from the water column over 2 hours in controlled laboratory experiments. It was hypothesised that oysters would filter amoebae from the water column, reducing the total water column amoeba load, and that amoeba DNA would accumulate in various oyster organs at different times, based on ingestion/egestion paths and proportionally to the total reduction in the water column.

## 2. Methods

### 2.1 Neoparamoeba perurans culture

A clonal strain of *N. perurans* described by (Botwright et al., 2020) was grown in a liquid malt yeast broth (MYB; 0.01 % (w/v) malt extract (Oxoid) and 0.01 % (w/v) yeast extract (Oxoid) in filtered, sterile seawater) at 15 °C (English et al., 2019) in 25 cm^2^ tissue culture flasks (Greiner Bio-One). Cultures were regularly tapped to remove adherence to the flask and subsequently split with new media weekly. On the day of the experiment, flasks of the *N. perurans* culture were vigorously tapped to dislodge the attached trophozoites, poured into a 50 mL sterile falcon tube, centrifuged (4000 x g, 5 min, 4 °C) to collect a pellet and resuspended in 2 mL MYB.

### 2.2 Experimental design

Oysters (*M. gigas*; 80–100 mm shell length) were obtained from Tasmanian Oyster Company (Tasmania, Australia) and held in a temperature-controlled room (17 °C) with fluorescent lighting (12:12 h day:night cycle) in 75 L of natural seawater. Oysters were fed daily with 15 mL Shellfish Diet 1800 (Reed Mariculture, Campbell, USA). Prior to the experiment, all oysters were depurated with daily 100 % water changes in artificial sea water (Aqua Forest Reef Salt, Buckinghamshire, United Kingdom; 35 ppt) and starved for 3 days to assist in the removal of transient bacteria, food particles, and potential DNA from their holding buckets. Twenty-four hours prior to the start of the experiment the oysters were placed in individual experimental buckets (n=9) with 2 L of artificial seawater and constant aeration. To ensure oysters were acclimated to the new buckets, visual observation to ensure an open valve was conducted, signalling respiration and/or filtering. Oysters that were not visually seen to have opened were not used for the experiment and were replaced with another replicate.

#### 2.2.1 Controls

Controls were taken throughout the experiment to ensure there was no contamination between any treatment, or background *N. perurans* signatures within the oyster organs. Buckets from all treatments, including experimental treatments (amoeba+water+oyster) and controls (amoeba+water, water only, oyster+water) were distributed within the room according to a random design. A water only (n=3) control consisted of artificial saltwater (ASW). Additionally, replicate buckets (oyster+water, n=3) to which *N. perurans* were not added were sampled as “negative, oyster” controls to address potential background signal within oyster organs. Positive controls (amoeba+water, n=3) consisted of *N. perurans*-inoculated buckets without oysters. Potential organ contamination from simply dipping an oyster in a tank with experimental quantities of *N. perurans* was also investigated (Time 0).

#### 2.2.2 Amoeba inoculation

Amoebae were enumerated in triplicates using a haemocytometer to determine the volume required to inoculate into each experimental tank to create a final concentration of 200 cells L^-1^. Due to the static inoculation concentration during the experiment, this quantity was chosen to ensure detectability by the end of the experiment. Additionally, it was chosen because it is higher than what is found in the environment, yet within the range seen in challenge models. AGD challenge models vary in their inoculation concentration, commonly ranging from 100 - 1000 cells L^-1^ (Adams et al., 2009, Downes et al., 2017) with some inoculating 10000 (Leef et al., 2005) to ensure rapid and reliable infection. At the beginning of each time course amoeba were inoculated into each 2 L tank.

#### 2.2.3 Video recording

All buckets were recorded for the entirety of the experiment using a GoPro Hero 11. This was done to quantify the duration of time oysters remained open while exposed to *N. perurans*. Upon completion of the experiment, recordings were viewed, and replicates in which the oyster was closed for >50 % of the time were removed from downstream analyses.

#### 2.2.4 Water samples

Nine experimental buckets (amoeba+water+oyster), 3 positive controls (amoeba+water) and 3 artificial salt water (ASW) controls (water only) were randomly sampled at each time point (0 h, 0.5 h, 1 h, 2 h post inoculation; Figure 1). The entire water sample (2 L) was filtered through a 0.22 µm mixed cellulose ester (MCE) disk filter membrane (Merck, New Jersey, USA) directly from buckets using a mechanical pump (300 rpm) immediately after oyster removal. Once filtered, membranes were removed from filter housings using sterile forceps and placed in a 2.2 mL microcentrifuge tube filled with 1 mL Longmire’s preservative solution (Longmire et al., 1997, Edmunds and Burrows, 2020).

**Figure 1:**
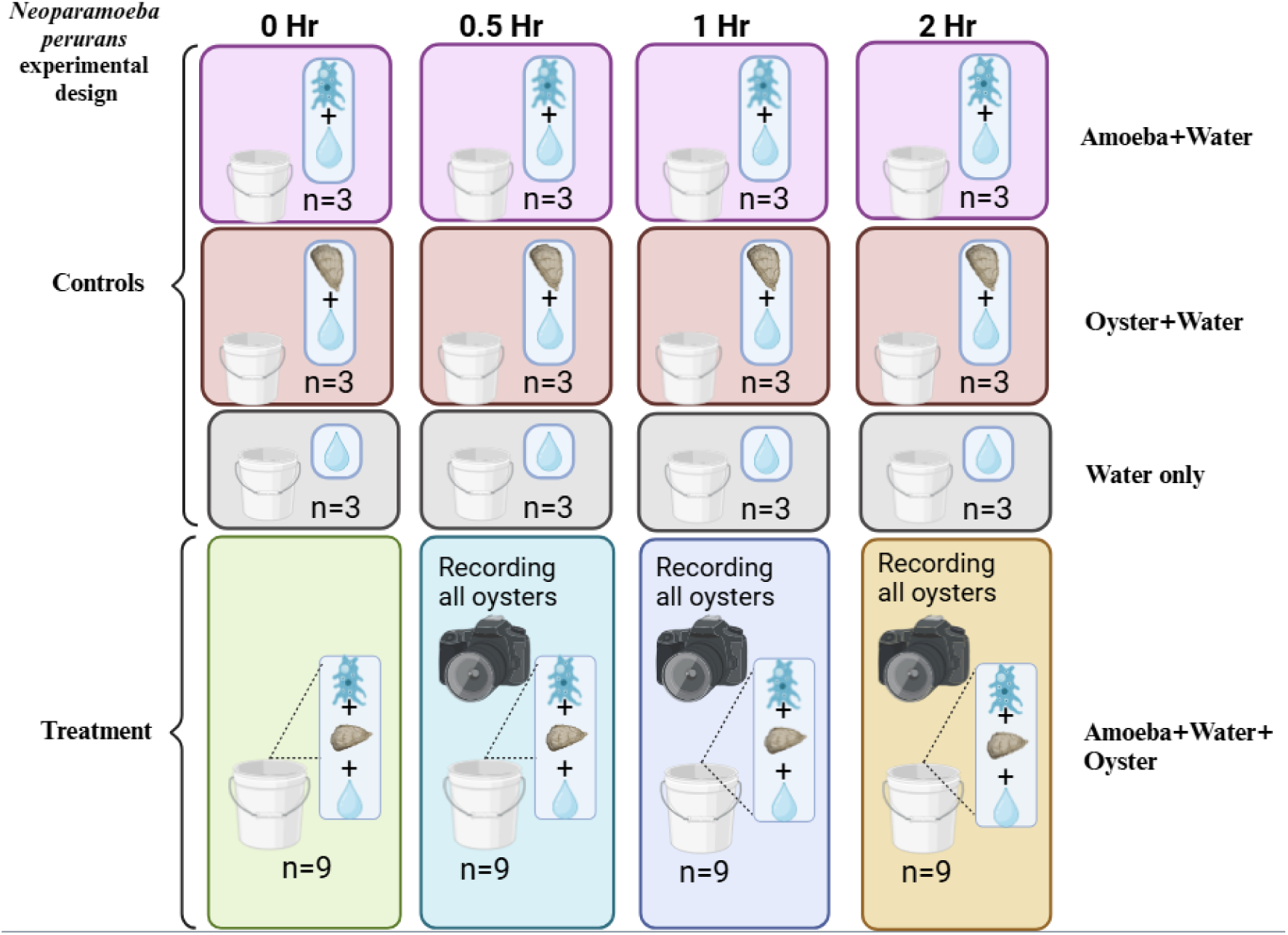
Experimental design for the inoculation of Neoparamoeba perurans into experimental buckets. Each time point consisted of the following controls: Oyster with no amoebae (purple, n=3), amoebae with no oyster (red, n=3), water with no oyster or amoebae (grey, n=3). At each time point (0, 0.5, 1, 2 h), nine replicate buckets (each containing a single oyster) were sampled.

#### 2.2.5 Oyster samples

At the defined time points oysters were removed from their holding buckets and briefly rinsed (by placing the oyster under constant flow for 5 s) with 0.22 µm filtered ultra-pure water (MilliQ), before being shucked with a sterile shucking knife. Sections of organs (∼1.0 g total wet weight; mantle, gill, palp, digestive gland) were removed using sterile forceps and scissors, with each individually placed in a sterile 2.2 mL microcentrifuge tube before being frozen (−80 °C). Once frozen, samples underwent a lyophilisation process at – 80 °C for 48 h. Lyophilised samples were homogenized using lysing matrix S tubes (MP Biomedicals, California, USA), using the FastPrep (MP Biomedicals, California, USA) homogeniser under the general program (6m s^-1^, 30 s) until powdered. Powdered organ samples were resuspended in 1 mL of Longmire’s preservative.

#### 2.2.6 Pseudofaece collection

Pseudofaeces were collected from buckets in which they were observed. These were identified by the loose clumping nature typically seen in pseudofaeces when compared to faeces and were found on the left side of the oyster, near the pseudofaece rejection region (Supplementary materials, S1) as described by Wisely & Reid (1978). If present, pseudofaeces were carefully removed using a sterile 1 mL pipette and placed in a sterile 2.2 mL microcentrifuge tube. These samples were then inspected under a microscope for visual detection of amoeba presence before being placed in 1 mL Longmire’s preservative buffer.

### 2.3 DNA extraction

DNA was extracted from organs, pseudofaeces, and from MCE filters stored in Longmire’s buffer, following the preserve, precipitate, lyse, precipitate, purify (PPLPP) protocol described by Edmunds and Burrows (2020), with minor modifications. One modification was the addition of 10 μL Proteinase K to each sample during lysis to reduce the digestion time to 2 h. To remove inhibitors all samples were additionally treated with the Zymo One-Step PCR Inhibitor Removal Kit (Zymo, California, USA) according to the manufacturer’s protocol.

### 2.4 Real-Time Quantitative PCR

#### 2.4.1 Neoparamoeba perurans

To quantify the copy number of the 18S rRNA gene of *N. perurans* within a sample, quantitative PCR (qPCR) was performed using a QuantStudio 5 Real Time PCR machine (ThermoFisher, Pennsylvania, USA). The total amoeba load (cells) was calculated by dividing the total 18S rRNA gene copies found by the number of copies of the 18S rRNA gene found within an amoeba (2880), as described by Bridle et al., (2010). Reactions were performed in triplicate, with primers specific to the *N. perurans* 18S rRNA gene, using previously described methods (Downes et al., 2015) with a modified annealing temperature. Each 10 µL reaction consisted of 5 µL PrimeTime Gene Expression Master Mix (Integrated DNA Technologies, Iowa, USA), 300 nM ‘NP1’ (5’-AAAAGACCATGCGATTCGTAAAGT-3’), 900 nM ‘NP2’ (5’-CATTCTTTTCGGAGAGTGGAAATT-3’), 200 nM ‘NPP’ (6-FAM-ATCATGATTCACCATATGTT-MGB), 2 µL template, and 0.8 µL DNA free water. Reactions then underwent the following cycling conditions: an initial denaturization of 95 °C for 2 min, followed by 45 cycles of 95 °C for 1 s and 60 °C for 20 s (modified from 56 °C). To create a standard curve based on total 18S rRNA gene copies, the Downes et al. (2015) amplicon was cloned into the pGEMT-easy plasmid (Promega) and transformed into TOP10 chemically component *E. coli* (Invitrogen, USA), as described by English et al. (2019).

Blue/white colour screened colonies with plasmid inserts (n=5) were miniprep purified and screened using the T7, Sp6 PCR primer pair. Plasmids were prepped for sequencing using the BigDye Terminator v3.1 Cycle Sequencing Kit before being purified with Agencourt ClenSEQ as per the manufacturers protocol. Cleaned plasmids were then Sanger sequenced (AGRF, Brisbane, Australia) to confirm the correct insert. All sequences matched the expected *N. perurans* sequence. Plasmid DNA was quantified using Qubit (Broad Range dsDNA kit, Thermo Fisher Scientific, Pennsylvania, USA) and copy number calculated based on plasmid size and DNA concentration. 1 - 10x serial dilutions of the plasmid (8.32 x 10^8^ down to 8.32 x 10^3^) were generated to create a standard curve. Standard curve efficiencies were within the range of 97 – 110 % and were calculated through the QuantStudio Design and Analysis 2.6.0 software.

#### 2.4.2 Beta-actin (*β*-actin)

Beta actin was used as a normalising gene, an internal standard for oyster tissue/DNA extracted and subsequently added to each qPCR reaction. Dry weight experiments were conducted to validate the internal standard. Gill samples (n=5) were removed from oysters and freeze dried before being homogenised and weighed out into four experimental weights (5 mg, 10 mg, 20 mg and 40 mg, Supplementary S2; Table S2). Each 10 µL reaction contained 5 µL SYBR Green (Bioline, Tennessee, USA), 400nM forward (5’-GTGCTACGTTGCCCTGGACT’-3’) and 400nM reverse primers (5’-TCGCTCGTTGCCAATGGTGA’-3’), 2 µL template DNA and 2 µL MilliQ water. Reactions underwent the following thermocycling conditions; initial hold at 95 °C for 2 min, then 40 cycles of 95 °C for 5 s, 60 °C for 30 sec, followed by a standard melt curve: 95 °C for 15 s, 60 °C for 1 min, 95 °C for 1 s (Du et al., 2013). To create standard curves, gBlock synthetic DNA standards (Integrated DNA Technologies, Iowa, USA) were created using the sequence obtained from the PCR product for Beta-actin. Once obtained, the dried pellet was resuspended to 10ng µL^-1^ stock. Copy numbers were calculated by the equation (copies= 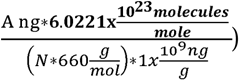 where *A*= amount of the amplicon produced (ng) and N= length of the amplicon. Once copy number was calculated, serial dilutions were completed (10^9^ copies µL^-1^ to 1 copy µL^-1^) with premade IDTE buffer (10 mM Tris, 0.1 mM EDTA) containing 0.5 mg mL^-1^ tRNA carrier to assist in the preservation of low copy number dilutions over extended periods of storage.

The mean of the technical replicates for both assays (Beta-actin and *N. perurans*) was only accepted if the standard deviation for the technical replicates was less than 25%. Samples that did not pass were run again until their SD was within the acceptable range. The melt curve for beta-actin was checked to ensure the amplification from each of the samples without dimer and non-specific binding.

### 3.0 Statistical analysis

A non-parametric one way Aligned Ranks Transformation Analysis of Variance (ART ANOVA) was conducted to test for differences in amoeba DNA quantity within the water column over time. Additionally, a separate model ART ANOVA was conducted comparing the positive controls over time. A factorial ART ANOVA was conducted with the ratio of amoeba/β-actin copies to test for differences in ratio in organ types (4 levels), over times (4 levels), and their interaction. A post-hoc pairwise comparisons test was conducted after the ART ANOVA to identify where significance lies within the dataset. Adjustments for multiple comparisons were made using the Tukey method for multiple comparisons. These models were chosen based on violations of the ANOVA assumptions, which were identified by conducting a Shapiro-Wilk Test for normality and a Levene’s Test to test for homogeneity of variances. Data transformations including square root, cube root and log10 +1 were evaluated to attempt to overcome both assumption violations, however, were unsuccessful. All statistics were conducted using R (v.4.2.2), and packages ARTool (Wobbrock et al.) for ART ANOVA and car (Fox, 2015) for normality testing.

## 4.0 Results

### 4.1 Assessment of oyster filtration

Analysis of video recordings showed that over 88 % of the oysters remained open for greater than 50% of the experimental period (2 h). Oysters that did not meet this threshold were removed from the analysis, resulting in removing two replicates at times 1 h and 2 h post amoebae inoculation (4 in total).

### 4.2 Depletion of amoeba from the water column

Amoebae were only detected in buckets into which they were inoculated. In buckets without oysters to which amoebae were added (positive control), there was no significant change in amoeba counts over time (*df*=3, *F*=2.8058, p>0.05). Total amoeba load decreased significantly from the water column over time in the presence of oysters (*df*=3, *F*=4.9355, p<0.01; Figure 2). At each time point we observed considerable variation in amoeba load across replicate water samples from buckets with oysters present. Results from the post-hoc test showed significant changes within the experimental buckets from time 0 h to 1 h (*df*=33, *T*=3.633, p<0.05) and 0 h to 2 h (*df*=33, *T*=4.201, p<0.01) post-inoculation of amoeba.

**Figure 2:**
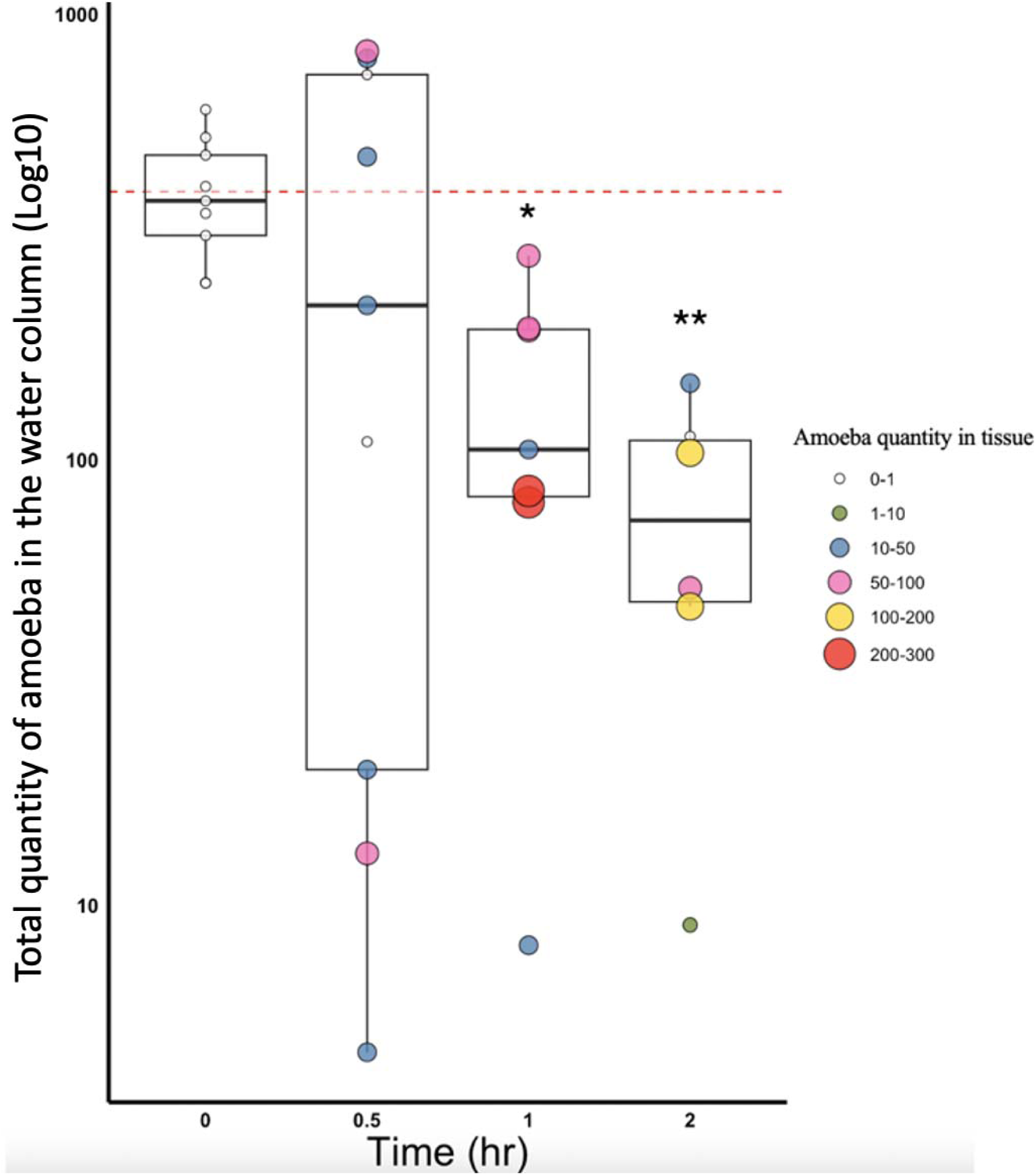
**Quantitative assessment of amoeba load in water samples and oyster tissues over time**. The boxplot shows amoeba counts (qPCR) in the water columns of experimental buckets over the course of 2 hours Replicate points indicate the total amoeba load found within the sum of the four organs (mantle, gill, palp, digestive gland), to show the relationship between amoeba loads in the water column vs oysters. Size and colour of each replicate indicate amoebae load found within the replicate. Red dashed line indicates amoeba load added to buckets at time 0 h. Significant differences from time 0 h are indicated by * where p<0.05 and ** where p<0.01.

There was a significant interaction effect of Time and Organ on amoeba counts in oyster tissue (*df*=9, *F*=2.6117, p<0.01), along with significant main effects of Time (*df*=3, *F*=23.6228, p<0.001) and Organ (*df*=3, *F*=11.2376, p<0.001). We hypothesized that different organs would accumulate amoeba at different times, so investigated differences in amoeba counts between organs within times and found a significant difference between the digestive gland and the palp at 1 h (*df*= 23, *T*=3.215, p<0.05). When we investigated the effect of time within an organ, we found significant differences in accumulation within the digestive gland from time 0 h to 1 h (*df*=27, *T*=-3.636, p<0.01). We also found significant differences within the palp at times 0 h to 2 h (*df*=27, *T*=-3.529, p<0.01) and 0.5 h to 2 h (*df*=27, *T*=-3.529, p<0.01; Figure 3). There were no significant differences within the gills or mantle between time points.

**Figure 3:**
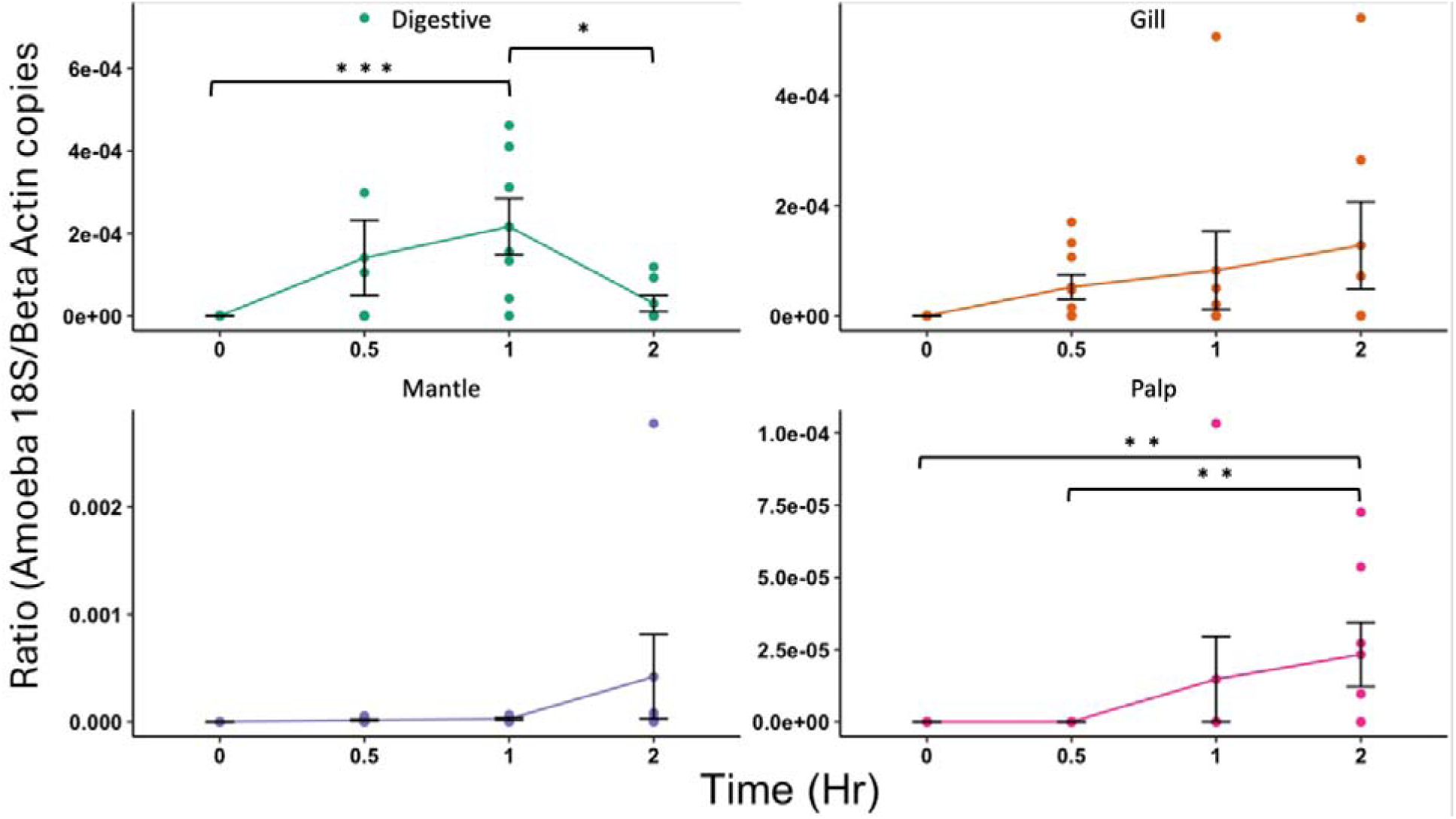
Line graph with error bars representing Amoeba 18S/ β-actin copy ratio calculated from the mean of three technical replicates obtained from the qPCR for four organs tested (Digestive, Gill, Mantle, Palp) over the course of 2 h post inoculation of amoeba. Error bars indicate SEM. Significant differences are indicated by * where p<0.05, ** where p<0.01 and *** where p<0.01.

### 4.3 Accumulation of amoeba within oyster organs

When evaluating results across all organs (Figure 4), amoebae were detected in over 88 % (8/9 replicates) of the oysters at time 0.5 h, 100 % (7/7 replicates) of the oysters at time 1 h, and 71 % (5/7 replicates) of the oysters at 2 h. The organ that had the most consistent detection was the digestive gland. The 1 h and 2 h timepoints had *N. perurans* detection in every organ (but not in every individual; Figure 4).

**Figure 4:**
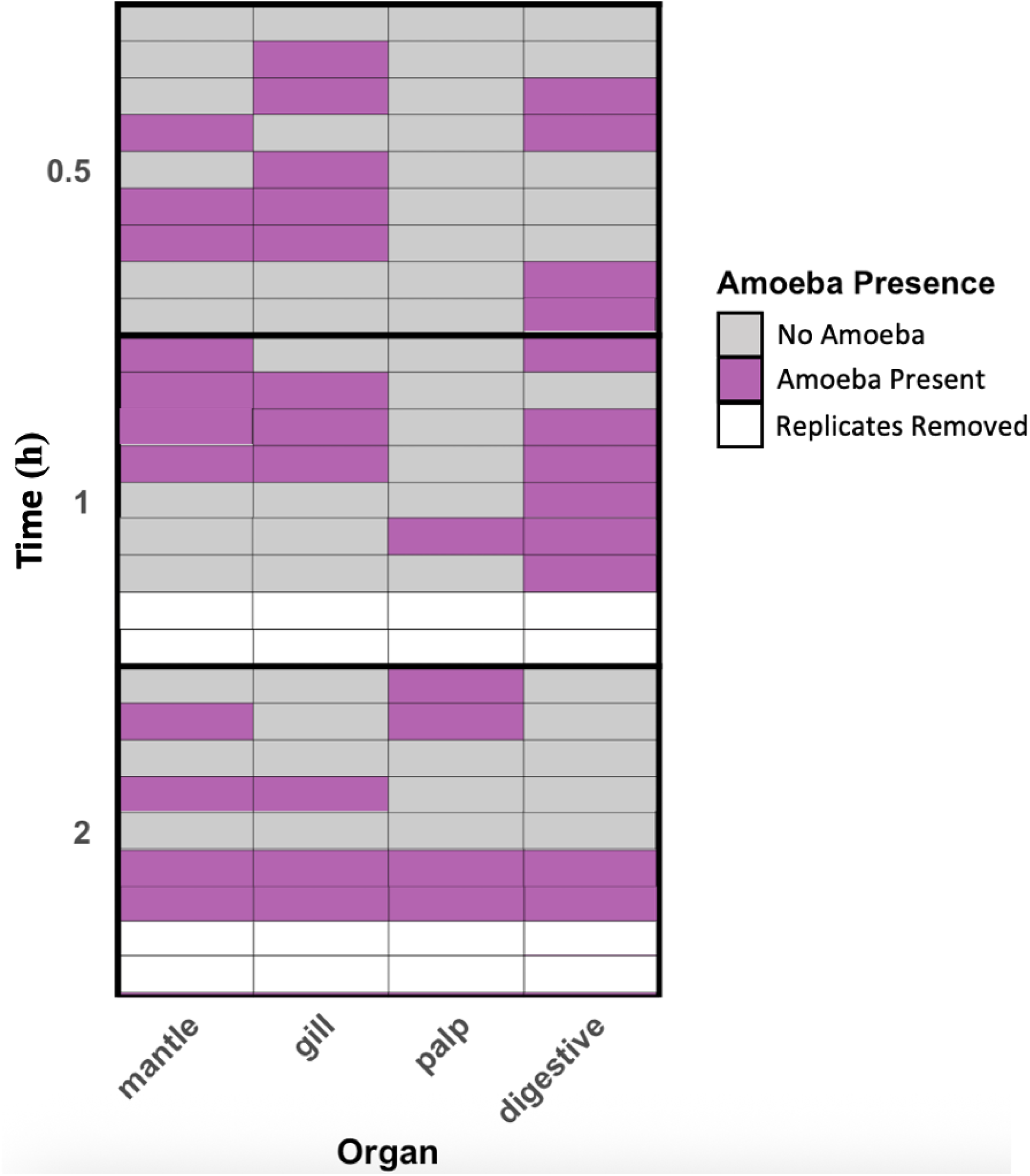
Amoeba presence/absence within organs at each experimental time point. Tile plot representing presence/absence of amoebae in each organ replicate at each time point (0.5 h, 1 h, 2 h). No amoeba DNA was detected at Time 0 h, therefore it is not shown. Grey tiles indicate the absence of amoeba, with purple indicating presence of amoeba. Each row indicates an individual. White tiles indicate the replicates removed from all analyses due to being closed for >50 % of the experimental time.

### 4.4 Amoeba detection within pseudofaeces

Pseudofaeces were not found in all buckets, including no visible pseudofaeces in control (oyster+water) buckets. They were, however, visible in 4 buckets at time 0.5 h, 4 buckets at 1 h and 3 buckets at 2 h post inoculation of amoeba. A subset of pseudofaece samples (n=3) showed intact amoebae when visualised under a microscope (Supplementary materials, S3). Despite this, detection via qPCR was unreliable. We were unable to achieve a SD below 25 % for technical replicates with many replicates having no amplification (Supplementary materials, Table S1).

## 5. Discussion

Natural samplers have previously been shown to provide a simple alternative to traditional eDNA sampling methods (Drinkwater et al., 2021). Natural samplers are cost-effective and easier to collect, enabling increased sample intensity. As described above, the use of bivalves as natural samplers addresses many of the attributes deemed necessary for a successful eDNA sampling campaign, including those targeting *N. perurans* given bivalves occur commonly in areas near fin fish farms, are easily collected, and can filter large quantities of water. This has great implications for Atlantic salmon aquaculture where monitoring and treatment for AGD contribute to nearly 20 % of the production costs of marine farmed salmon (Munday et al., 2001). Due to the need to continuously treat affected cohorts to minimise AGD related production losses, detection of *N. perurans* has been the focus of many AGD studies (Bridle et al., 2010, Wright et al., 2015, Wright et al., 2017, Taylor et al., 2021, Wynne et al., 2024). The use of molecular methods has shown sensitive and reliable detection of *N. perurans* from the water column through pump-based filtration methods. In this study, we aimed to explore the use of *Magallana gigas* as a natural sampler for accumulation of *N. perurans* DNA.

This study provides the first evidence that *N. perurans* can be detected reliably within oyster organs. The greatest number of detections of amoeba was found within the digestive gland, demonstrating oysters were ingesting amoebae, a novel finding as no literature has reported this. With Pacific oysters being common in marine environments, including near fish pens, reliable detection of *N. perurans* within any tissue provides a potentially simplified method for *N. perurans* detection on finfish aquaculture leases.

We show that qPCR-based detection of amoeba is possible from Pacific oysters exposed to experimental tanks containing 200 amoeba L^-1^. A concentration of amoeba that has been shown to cause AGD in challenge trials (Adams et al., 2009; Downes et al., 2017). This contrasts with earlier work by Rolin and colleagues where no detection was found in blue mussels was demonstrated using a concentration of 3000 amoeba L^-1^, indicating that utilising *M. gigas* as a sampler provides greater sensitivity and probability of detection than the blue mussel.

The amoeba load within the water column significant decreased over 2 h. While there may have been adherence of amoeba to experimental surfaces, as seen in previous studies on mussels (Rolin et al., 2016), this was not the primary cause of reduction in the water column as no decline was observed in our amoeba+water controls and the concomitant detection of amoeba DNA within oyster tissues suggest the presence of oysters was the primary driver of amoeba concentration decline. Attachment to oyster shells was not directly assessed in the current study, but it is possible that some could be attributed to this mechanism.

Detection within various tissues throughout the time course offers evidence that oysters ingested amoebae and accumulated them within their organs, resulting in their reduction from the water column. However, we found no relationship between the numbers of amoeba removed from the water column and numbers accumulating within oyster organs. We assumed that there would be homogeneity of DNA distribution across the organ that was sampled. However, patchy distribution could account for some amoeba not being detected within the sampled subset of organs (∼1 g wet weight). While we expected that accumulation would reach a steady state within the organs, given there was constant exposure to amoeba for the duration of the experiment, this was not seen. This could be explained by quick digestion of DNA when in contact with digestive fluid (∼10 min) as described by (Bernard, 1974), or by the attachment of amoeba to bivalve shells as described by Rolin and colleagues (2016). While we measured whether the oysters were open/closed, this does not give an indication as to the rate of filtration as oysters open valves for more than just feeding (Famme, 1980) and could be a focus for future work.

Nearly all oysters exposed to amoebae tested positive in at least one organ, with the lowest detection rate (71% of replicates) occurring at 2 h post-inoculation, exceeding the highest detection (11%) reported in blue mussels on an Atlantic salmon farm (Hellebø et al., 2017). However, this could also be due to the controlled experiments used in this trial rather than *in situ* conditions used in the previous study and with much lower concentrations commonly found in the environment. While controlled experiments are advantageous in testing single variables, oyster feeding selectivity may change when exposed to a complex matrix seen in natural environments. Thus, oyster exposure to various concentrations of amoeba and integration of amoeba into a complex diet should be explored further to confirm whether the results in this controlled experiment compliment those found in environmental situations.

Amoebae were expected to be detected first in the mantle and gills, which initially accumulate food particles before sorting by the palps (Kellogg, 1915). As water and particles only briefly contact the mantle frills prior to being moved into the pallial cavity, concentrations within it are likely more dilute than in other organs. Unlike Rolin et al. (2016), who pooled mantle and gill samples, our data indicate the mantle is the least reliable organ for *N. perurans* detection. We recommend pooling the gill and digestive gland instead, as they yielded the most consistent detections across all time points. Similarly, Hellebø et al. (2016) reported sporadic, but positive detections in gill and digestive gland tissue of *Mytilus* spp. While individual organ results varied, potentially due to heterogeneous clumping of DNA across the organ, nearly all replicates tested positive when assessed at the whole-organism level, suggesting pooling may improve reliability. Dissection of individual organs can be time consuming and thus a pooled sample may increase ease of use.

Unwanted particles, or those in high abundance, initiate a rejection response and pseudofaeces are expelled from the pallial cavity (Loosanoff and Engle, 1947, Wisely and Reid, 1978). Pseudofaeces production was quite variable within this experiment.

Pseudofaeces were collected in all tanks in which they were seen (n=11 out of 23). Of those, three samples were randomly selected, and amoeba were confirmed to be visually present. This was initially done to compliment qPCR findings of amoeba detections, though qPCR detection of amoeba within pseudofaeces were unreliable. Where amoebae were visually detected in pseudofaeces, they were also found within the digestive glands of the same individuals, indicating that their presence did not initiate a total rejection response. While we did not test for the viability of amoeba within pseudofaeces, future work could aim to address this and determine reduction of amoeba load in the context of bioremediation.

Our results show that the natural sampling approach is an effective method to collect DNA of *N. perurans* from the water column, however there are caveats associated with its use. Quantity of amoeba within the water column is poorly studied, with most of the work focusing on the detection of amoeba rather than quantification. A previous study that quantified amoeba loads within the water column found concentrations from 0-62 amoeba L^-1^(Wright et al., 2015). While our study inoculated 200 amoeba L^-1^, we did not test limits of detection, how long amoeba remain within the oyster, or how much the oyster filtered. However, any detection within oysters offers evidence of the presence of amoeba within the water column.

Detecting amoebae, even at low concentrations, may assist in predicting AGD onset. In challenge models, onset of disease can be caused from extremely low quantities of amoeba (0.1 cells L^-1^) (Bridle et al., 2021) signalling that any presence of amoeba within a system may be indicative of AGD onset. Thus, presence/absence data for *N. perurans* is likely an important measure for informing risk to stock. There is only a single published challenge trial associating the load of amoeba with disease onset in salmon farming (Bridle et al., 2021), more research is needed, both in challenge trials and *in situ*. Future research could explore this to identify the relationship between pathogen load within the water column, level of amoebae detection in oyster organs, and disease onset/progression within salmon aquaculture.

## Conclusions

Our study has demonstrated the use of Pacific oysters, *M. gigas*, as a natural sampler for the detection of *Neoparamoeba perurans.* To our knowledge, this is the first time *N. perurans* has been consistently detected within organs of any bivalve species, and the first evidence of *M. gigas* ingesting this amoeba. Like mussels, oysters occur commonly in the aquatic environment and near Atlantic salmon farms. Therefore, these results suggest oysters could provide a simplified method to detect amoeba presence in a system. Thus, we show that the natural sampling approach using oysters potentially offers a straightforward method of collecting eDNA for *N. perurans* detection.

## Permits

All work was done under the POMS Group Movement permit and POMS 23-990.

## Data availability statement

Data pertaining to this work can be doing at doi.org/10.25919/96h1-ra18

## Acknowledgements

The authors acknowledge the financial support of the Blue Economy Cooperative Research Centre, established and supported under the Australian Government’s Cooperative Research Centres Program, grant number CRC-20180101. Additionally, we would like to thank Oysters Tasmania, with specific mention of Francis Huddlestone, for facilitating the sourcing of oysters as well as Tasmanian Oyster Company for supplying the oysters for the experiment. We acknowledge our other industry partner Petuna for their financial contributions to the work. The authors would like to thank Richard Taylor for his assistance in culturing the amoeba used in these experiments.

## Supplementary materials

**S1:**
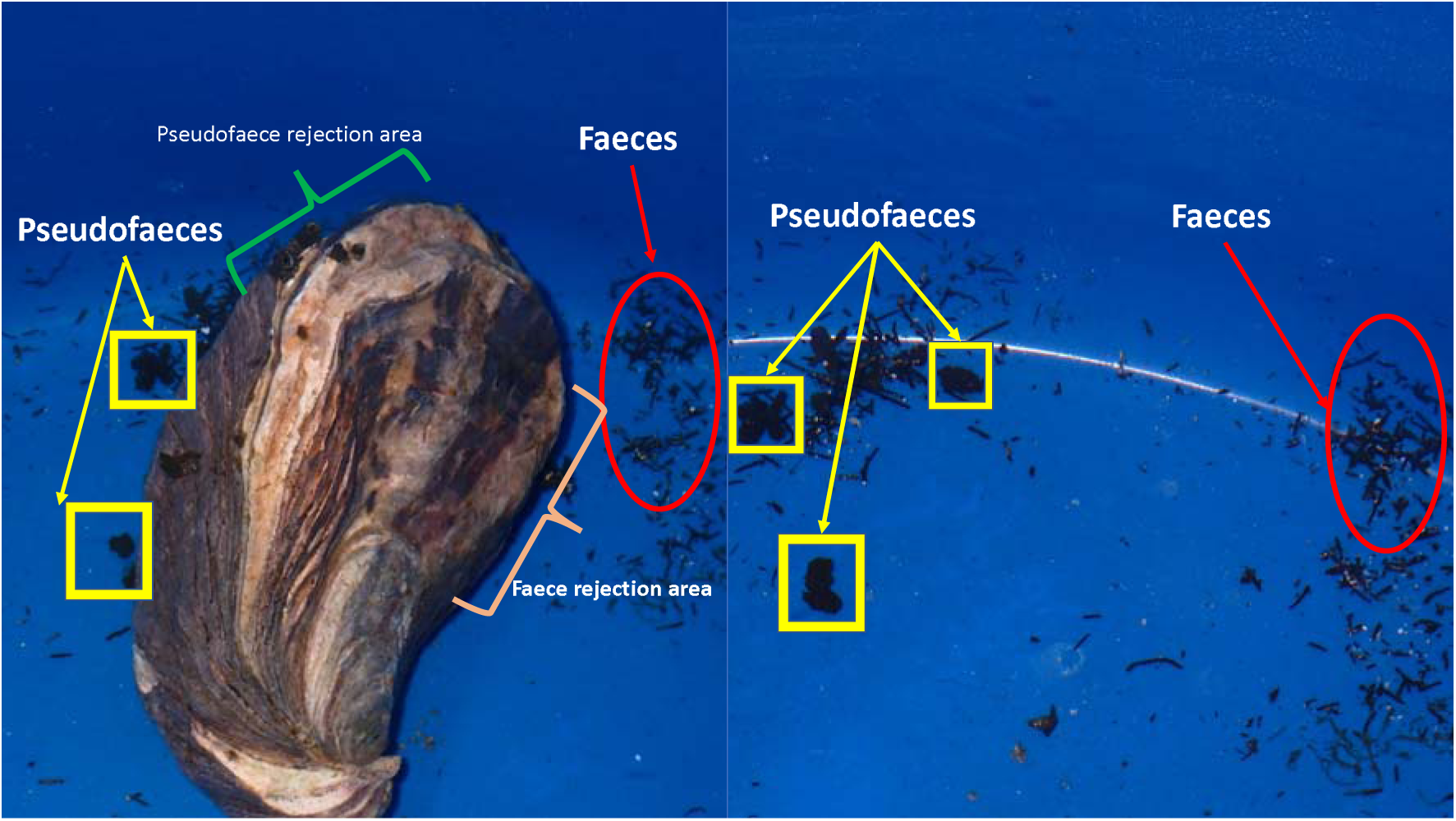
Images of the pseudofaece rejection area (green) and faece rejection area (orange) of the Pacific oyster (Magallana gigas). Pseudofaeces can be seen boxed in yellow and are loosely bound together and have no definite shape. Faeces are circled in red and have pellet forms in a defined shape.

**Figure S3:**
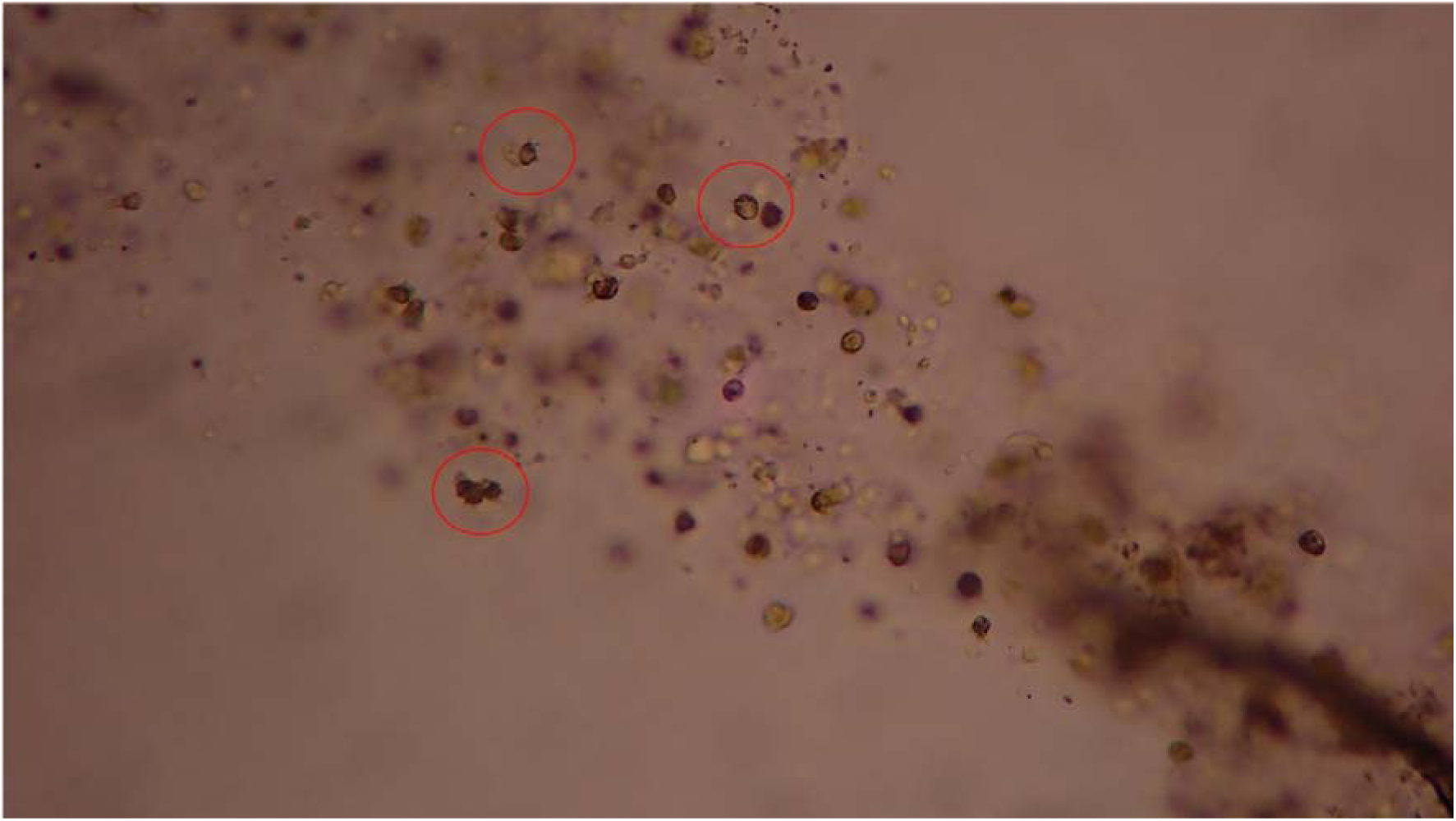
Pseudofaeces under the microscope. Red circles indicate where amoebae are entrapped in the mucous of the pseudofaeces. Amoeba were identified by their visibly extended pseudopodia.

**Table S1:** Pseudofaece qPCR results. Each tank represents replicates that had pseudofaeces visually observed and were extracted. Technical qPCR replicates and their positive amplification out of the total are shown for each respective sample.

| **Tank**<br>**(Replicate)** | **Time**<br>**(h)** | **Technical replicate positive**<br>**amplification** |
| --- | --- | --- |
| Tank 35 | 0.5 | 2/3 |
| Tank 36 | 0.5 | 1/3 |
| Tank 30 | 0.5 | 3/3 |
| Tank 31 | 0.5 | 1/3 |
| Tank 50 | 1 | 3/3 |
| Tank 51 | 1 | 2/3 |
| Tank 52 | 1 | 3/3 |
| Tank 54 | 1 | 1/3 |
| Tank 18 | 2 | 0/3 |
| Tank 13 | 2 | 1/3 |
| Tank 12 | 2 | 1/3 |

